# The functional specialization of visual cortex emerges from training parallel pathways with self-supervised predictive learning

**DOI:** 10.1101/2021.06.18.448989

**Authors:** Shahab Bakhtiari, Patrick Mineault, Tim Lillicrap, Christopher C. Pack, Blake A. Richards

## Abstract

The visual system of mammals is comprised of parallel, hierarchical specialized pathways. Different pathways are specialized in so far as they use representations that are more suitable for supporting specific downstream behaviours. In particular, the clearest example is the specialization of the ventral (“what”) and dorsal (“where”) pathways of the visual cortex. These two pathways support behaviours related to visual recognition and movement, respectively. To-date, deep neural networks have mostly been used as models of the ventral, recognition pathway. However, it is unknown whether both pathways can be modelled with a single deep ANN. Here, we ask whether a single model with a single loss function can capture the properties of both the ventral and the dorsal pathways. We explore this question using data from mice, who like other mammals, have specialized pathways that appear to support recognition and movement behaviours. We show that when we train a deep neural network architecture with two parallel pathways using a self-supervised predictive loss function, we can outperform other models in fitting mouse visual cortex. Moreover, we can model both the dorsal and ventral pathways. These results demonstrate that a self-supervised predictive learning approach applied to parallel pathway architectures can account for some of the functional specialization seen in mammalian visual systems.

## 1 Introduction

In the mammalian visual cortex information is processed in a hierarchical manner using two specialized pathways [15, 52]: the ventral, or “where” pathway, and the dorsal, or “what” pathway. These two pathways are specialized for visual recognition and movement, respectively [42, 24, 19, 58, 59, 17]. For example, damage to the ventral pathway may impair object recognition, whereas damage to the dorsal pathway may impair motion perception [64].

Deep artificial neural networks (ANNs) trained in a supervised manner on object categorization have been successful at matching the representations of the ventral visual stream [62, 57, 32]. They have been shown to develop representations that map onto the ventral hierarchy, and which can be used to predict [62] and control [1, 49] neural activity in the ventral pathway. However, when we look at the other principal visual pathway in the mammalian brain, i.e. the dorsal pathway, the situation is different. Very few studies have examined the ability of deep ANNs to develop representations that match the dorsal hierarchy (though see the following fMRI study: [21]). Moreover, to the best of our knowledge, no studies have demonstrated both ventral-like and dorsal-like representations in a single network.

This lack of deep ANN models that capture both ventral and dorsal pathways leads naturally to an important question: under what training conditions would specialized ventral-like and dorsal-like pathways emerge in a deep ANN? Would a second loss function be required to obtain matches to dorsal pathways, or is there a single loss function that could induce both types of representations?

One promising set of candidates are predictive self-supervised loss functions [45, 22, 38]. Recent work has shown that self-supervised learning can produce similar results to supervised learning for the ventral pathway [63, 34, 26]. Moreover, there is a large body of work showing that mammals possess predictive processing mechanisms in their cortex [31, 8, 30, 18, 16], including in the dorsal pathway [2, 35], which suggests that a predictive form of self-supervised learning could potentially lead to the emergence of both ventral and dorsal-like representations.

Addressing this question requires recordings from different ventral and dorsal visual areas in the brain. Here, we explore these issues using publicly available data from the Allen Brain Observatory [9], which provides recordings from a large number of areas in mouse visual cortex. We examine the ability of a self-supervised predictive loss function (contrastive predictive coding [45, 22]) to induce representations that match mouse visual cortex. When we train a network with a single pathway, we find that it possesses more ventral-like representations. However, when we train a network with two parallel pathways we find that the predictive loss function induces distinct representations that map onto the ventral/dorsal division. This allows the network to better support both object categorization and motion recognition downstream tasks via the respective specialized pathways. In contrast, supervised training with an action categorization task only leads to matches to the ventral pathway, and not the dorsal pathway.

Altogether, this work demonstrates that the two specialized pathways of visual cortex can be modelled with the same ANN if a self-supervised predictive loss function is applied to an architecture with parallel pathways. This suggests that self-supervised predictive loss functions may hold great promise for explaining the functional properties of mammalian visual cortex.

## 2 Background and related work

### Self-supervised ANN models of the ventral visual stream

Recently, [63] and [34] showed that the representations learned by self-supervised models trained on static images produce good matches to the ventral pathway. Our work builds on this by exploring the potential for self-supervised learning to also explain the dorsal pathway.

### ANN models of the mouse visual system

Mice have become a common animal model in visual neuroscience due to the sophisticated array of experimental tools [44, 27]. As such, [53] compared the responses of different areas in mouse visual cortex with the representations of a VGG16 trained on ImageNet. In this paper, we show that self-supervised learning can produce better fits to both ventral and dorsal areas than supervised learning.

### ANN models of the dorsal pathway

The MotionNet ANN model [51], is a feedforward ANN trained to predict the motion direction of segments of natural images, which was inspired by the role of the dorsal pathway in motion perception. In [6], they show that learning both ventral and dorsal-like representations in a single ANN with two pathways is possible if one forces the two pathways to process the phase and amplitude of a complex decomposition of the stimuli separately. In [21], they show that training a deep ANN on supervised action recognition can induce some match to dorsal pathway fMRI recordings. In this study, we show that with prediction as the learning objective and an architecture that has two parallel pathways, both ventral-like and dorsal-like representations can be learned.

## 3 Methods

### 3.1 Datasets

We use the Allen Brain Observatory open 2-photon calcium imaging dataset for the experiments in this study. We select subsets of the dataset based on brain area, recording depth, and visual stimuli used. Recordings from five areas of mouse visual cortex are used (VISp, VISlm, VISal, VISpm, VISam; Figure 1a). We only exclude one area from our analyses (VISrl) because it is a multi-sensory area, and visual stimuli alone do not drive it well [9]. We use recordings from cortical depths of 175-250*μm* (which corresponds to cortical layers 2-3) as these are the recordings with the largest number of neurons. When selecting visual stimuli, we use recordings elicited by the presentation of natural movies, because unlike static images, movies can elicit clear responses in both ventral and dorsal areas [9]. Thus, we only use those parts of the dataset in which natural movies (30 seconds long) were presented as visual stimuli. More details about the dataset can be found in [9]. For training the deep ANNs, we use the UCF101 dataset [56] (see section A.4 of supplementary materials).

**Figure 1:**
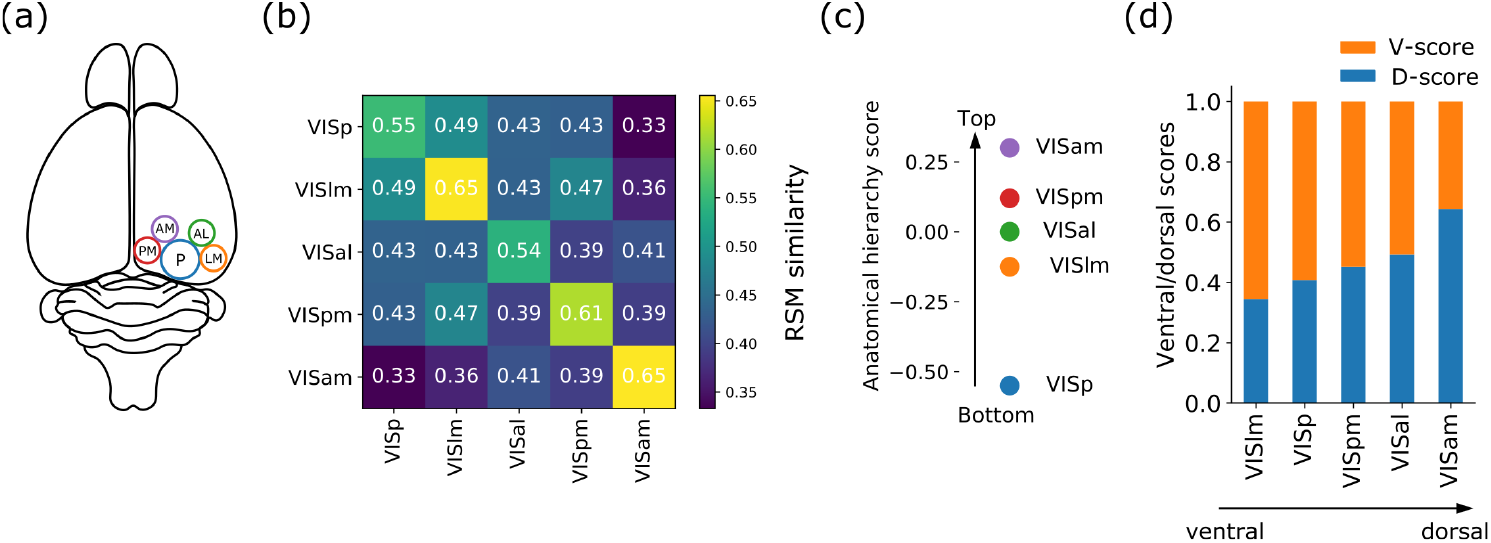
(a) The schematic of mouse visual cortex. (b) Representational similarity analysis between the visual areas included in our analysis. (c) The anatomical hierarchy score of the visual areas adopted from [23]. (d) Ventral and dorsal scores of the visual areas. Areas are sorted from the most ventral (VISlm - left) to the most dorsal (VISam - right) areas.

### 3.2 Analysis techniques

#### Representational Similarity Analysis (RSA)

To measure the representational similarities between real brains and ANNs we use the RSA method. RSA has been commonly used in both the neuroscience [11, 33] and deep learning literature [41, 43]. The details of RSA can be found in those citations, but we will summarize it briefly here. First, we create response matrices 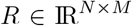 for every brain area and every layer of our ANNs, where *N* is the number of neurons and *M* is the number of video blocks, with each block comprising 15 frames of the video. Element *ij* of the response matrix represents the response of the *i^th^* neuron to the *j^th^* video sequence. We then use Pearson correlation to calculate the similarity of every pair of columns (*e.g. k^th^* and *l^th^* columns) in matrix *R*, and form the *M* by *M* Representation Similarity Matrix 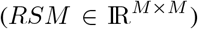, in which every element (*kl*) quantifies the similarity of the responses to the *k^th^* and *l^th^* videos blocks. Thus, the *RSM*s describe the representation space in a network, be it a brain area or an ANN layer. Given two *RSM*s, we then use Kendall’s τ between the vectorized *RSM*s to quantify the similarity between the two representations. It is important to note that measurement noise can potentially induce bias in the RSM estimations, but since the variance of measurement noise is not expected to be different across different video blocks, using Kendall’s *τ* rank correlation should cancel the bias in our RSM estimations (see [12] for more details). As an additional sanity check, we use RSA to compare the representations between mice. If RSA is identifying salient representational geometries, then the *RSM*s between different areas should be lower than the RSMs for the same areas. Indeed, as shown in Figure 1b, the representational similarity is highest for the same regions across animals (i.e. the diagonal values are larger). These diagonal values also represent the noise ceiling for these areas (see section A.1). Throughout the paper (with the exception of Figure 1b), *RSM* similarities are reported as the percentage of the noise ceiling.

#### Identification of brain regions in the hierarchy

The hierarchical organization of mouse visual areas can be inferred based on the anatomical and functional properties of each brain region. We adopted an approximation of hierarchical indices as reported in [23]. The relative hierarchical placing of the visual areas included in this study are shown in Figure 1c.

#### Identification of brain regions into ventral and dorsal streams

We group the brain regions into two sets, i.e. ventral and dorsal areas. Although the ventral/dorsal specialization of visual pathways in mice is not as clear and well understood as it is in primates, many anatomical and physiological studies do suggest that mice also possess such specialized pathways [40, 58, 39, 37]. According to previous anatomical and physiological studies [17, 58], VISlm and VISam can be considered as the most ventral and dorsal areas of mouse visual cortex, respectively. We then use the similarity of representations between other areas and VISlm and VISam to estimate a ventral score 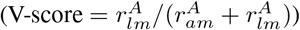 and a dorsal score 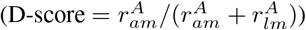 for each area. 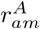 is the representational similarity between area *A* and VISam, and 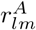 is the representational similarity between area *A* and VISlm. The D-score and V-score values for the five visual areas are plotted in Figure 1d. Based on the D-score and V-score values, we grouped VISlm, VISp, and VISpm as more ventral and VISal, and VISam as more dorsal in our analysis. This grouping is in keeping with a recent study that showed VISam and VISal are the first and second most responsive areas to motion stimuli: an important characteristic of dorsal areas [54].

### 3.3 Model

#### Self-supervised predictive learning

We used Contrastive Predictive Coding (CPC) for self-supervised learning, which was developed for modeling sequential data, including video datasets [45]. The loss function relies on predicting the future latent representations of a video sequence, given its present and past representations. See Figure S1 for a schematic of the model, and A.2 for more details regarding the CPC loss function.

#### ANN Architecture

All the ANN backbones used in our study are variants of the ResNet architecture, similar to the ones used in [13]. Our ResNet architectures have either one or two pathways. The one-pathway ResNet (ResNet-1p) is a regular 3D ResNet (Figure 2a). The two-pathway ResNet (ResNet-2p) is composed of two parallel ResNet branches, which split after a single convolutional layer, and merge after their final layers through concatenating their outputs along the channel dimension (Figure 3a). Both pathways of ResNet-2p receive a copy of the first convolutional layer output, and each has 10 Res-blocks. To keep the total number of output channels the same in both architectures, each pathway in ResNet-2p has half the number of channels of the single pathway in ResNet-1p. For the ANNs trained on object categorization, we use ResNet-18 pretrained on the ImageNet database [25]. Our 3D ResNet architectures are summarized in Table S1.

**Figure 2:**
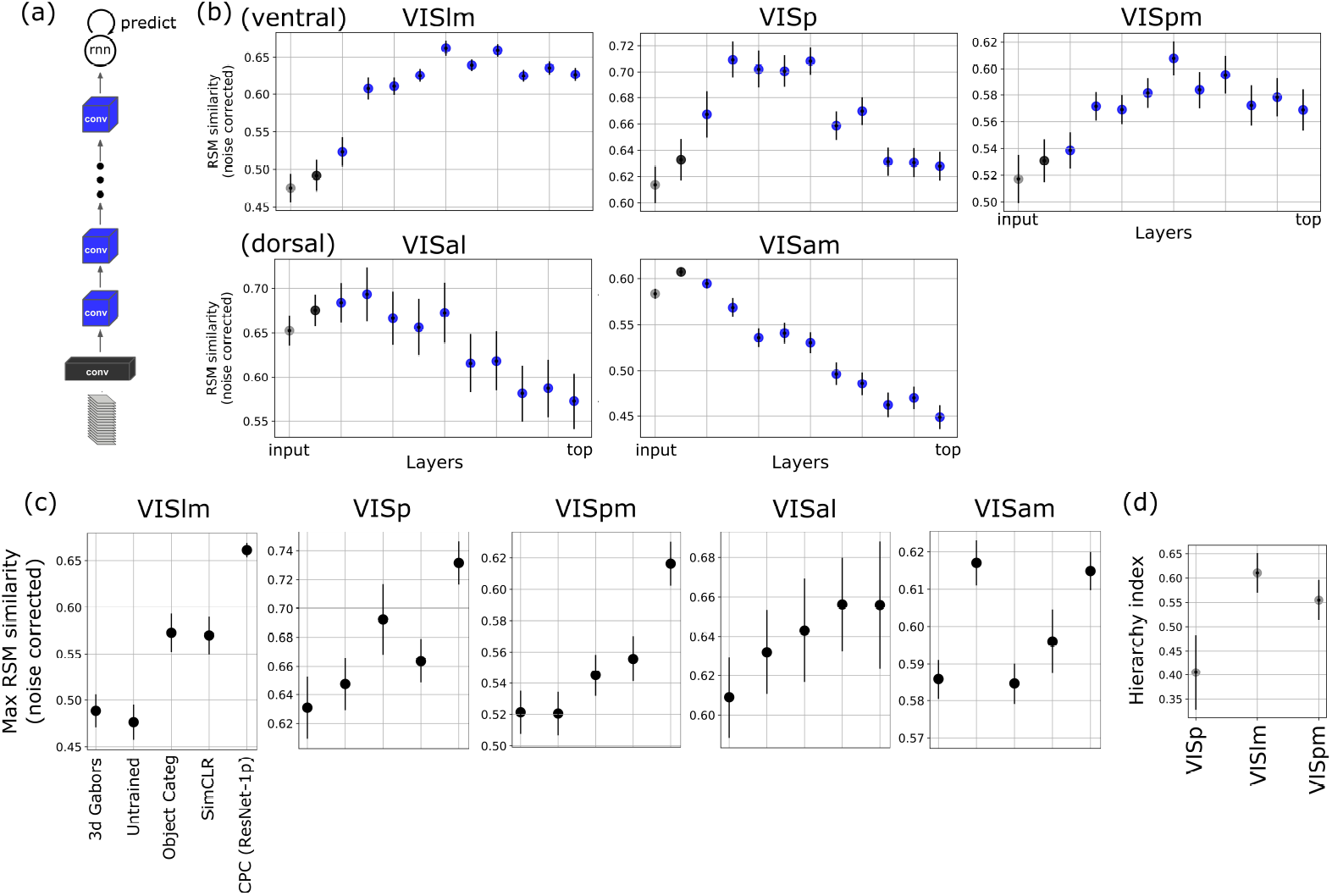
Representational Similarity Analysis between all the visual areas and the ANN trained with CPC. (a) The schematic of the ANN architecture with one pathway (ResNet-1p) used as the backbone of CPC. (b) Representational similarity between all the layers of the ANN with one pathways (trained with CPC) and the ventral (top: VISlm, VISp, VISpm) and the dorsal (bottom: VISal, VISam) areas. (c) The maximum representational similarity values between the ANN and the ventral and dorsal areas. (d) Hierarchy index of the ventral areas based on their fit to the ANN. Error bars represent bootstrapped standard deviation.

**Figure 3:**
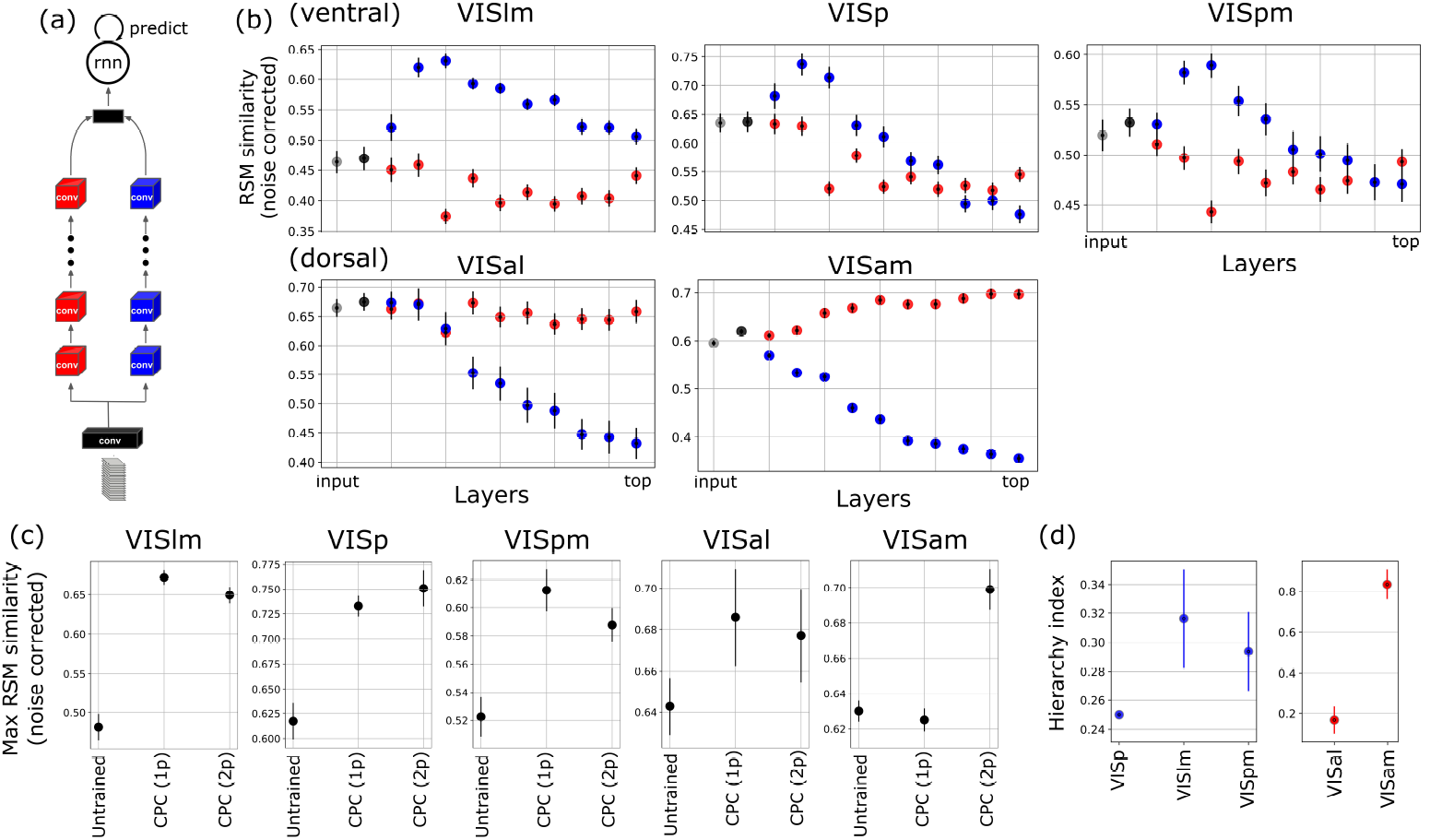
Representational Similarity Analysis between all the visual areas and the ANN trained with CPC. (a) The schematic of the ANN architecture with two pathways (ResNet-2p) used as the backbone of CPC. (b) Representational similarity between all the layers of the ANN with two pathways (trained with CPC) and the ventral (top: VISlm, VISp, VISpm) and the dorsal (bottom: VISal, VISam) areas. (c) The maximum representational similarity values between the ANN and the ventral and dorsal areas. CPC (1p) and CPC (2p) are ResNet-1p and ResNet-2p, respectively, both trained with CPC loss function.(d) Hierarchy index of the ventral (left; in red) and dorsal (right; in blue) areas based on their fit to the ANN. Error bars represent bootstrapped standard deviation.

#### Baseline models

In terms of alignment with brain representations of video sequences, we compare the CPC trained ANNs with four other models: (1) a simple model based on Gabor filters, (2) a randomized deep ANN, (3) a ResNet-18 trained on ImageNet, and (4) 3D ResNets trained on action recognition in a supervised manner. See section A.3 of supplementary materials for more details.

#### Training

We use the backpropagation algorithm and Adam optimizer. CPC is trained with a batch size of 40 samples, a learning rate of 10^−3^, and 100 epochs. Supervised action recognition is trained with a batch size of 256 samples, a learning rate of 5 × 10^−4^, and 300 epochs. All the models are implemented with PyTorch 1.7 and trained on RTX8000 NVIDIA GPUs.

#### Downstream tasks

In addition to comparisons with dorsal and ventral representations in mouse brain, we examine the two pathways of our trained ResNet-2p on two downstream tasks: object categorization and motion discrimination, which are supported in the real brain by the ventral and dorsal pathways, respectively. See section A.5 of supplementary materials for more details.

## 4 Results

### 4.1 CPC with a single pathway architecture produces better matches to mouse ventral stream

We first compare the representations learned with CPC on a single pathway architecture (Fig. 2a) with the three ventral areas of mouse visual cortex (VISlm, VISp, and VISpm; based on D-scores/V-scores in Figure 1d). In Figure 2b (top), the similarity of RSMs between different layers of the ANN and these three areas are shown. Comparing the maximum representational similarities between models in Figure 2c, we can see that, for the three ventral areas, CPC shows a higher maximum similarity compared to the other models. Compared to the baseline models (untrained ANN and Gabors), the ANN trained with object categorization has higher representational similarity to VISlm and VISp, the two most ventral areas (see Figure 1d), which is consistent with the suggested role of ventral areas in form and shape representation [10, 17]. This is in contrast with a previous study [5] that could not find any significant difference between ANNs trained on object categorization and untrained ANNs in modeling mouse visual cortex. There are several sources of variability that could explain this contradiction (e.g. architecture, datasets, etc.), but an important possibility is that the two studies used different visual stimuli, namely, natural videos in our study vs static natural images in [5]. Natural videos are better suited for eliciting sufficiently strong responses that can distinguish between the representations of the object categorization-trained and randomly initialized ANNs [9].

As noted, and as can be seen in Figure 2b, the maximum similarity happens in different hierarchical levels of the CPC-trained ANN for each area. We quantify the hierarchy index for every area by dividing the layer number with the highest similarity by the maximum number of layers. Based on this measure, the areas at the very top and bottom of the visual hierarchy have a hierarchy index of 1 and 0, respectively. The hierarchy index for the three areas are shown in Figure 2d. In accordance with the anatomical and functional data (Figure 1c), VISp has a lower hierarchical index (0.41 ± 0.15) than VISlm (0.61 ± 0.08) and VISpm (0.55 ± 0.08). It should be noted, that despite the fact that anatomical/functional hierarchy indices position VISpm higher in the hierarchy than VISlm, our model-based measure of hierarchy places the two areas at around a similar level, with VISlm having a marginally higher value than VISpm. We speculate that this may be a result of the fact that VISpm has noisier representations, and therefore a lower noise ceiling (Fig. 1b). This may be due to VISpm actually being more multi-modal than purely visual.

### 4.2 CPC on a single pathway does not match mouse dorsal stream

We compare the similarity of RSMs between ResNet-1p trained with CPC (Fig. 2a) and the dorsal areas (VISam and VISal; based on D-scores/V-scores in Figure 1d). As shown in Figure 2b (bottom), the ANN trained with CPC just passes the pixel-level representations in its early layers (the first gray circle in the plots), and then the similarity values continuously decrease for deeper layers. Notably, for the most dorsal area (VISam), the maximum similarity of RSMs does not go above the untrained model (Figure 2c). For VISal, even though CPC shows some improvement compared to an untrained ANN, the performance is much lower than for the ventral areas, as shown in the previous section (Fig. 2c). These findings show that the representations learned by CPC and the ResNet-1p architecture are more ventral-like, and do not easily explain dorsal area representations (see supplementary section E). Similarly, an object categorization trained ANN has a very low similarity with the dorsal areas (even lower than the untrained ANN for VISam) indicating that object categorization cannot be considered as an appropriate loss function for the dorsal pathway, which is consistent with our current understanding in neuroscience.

### 4.3 An architecture with parallel pathways trained with CPC can model both ventral and dorsal areas

CPC’s tendency to learn ventral-like representations, as seen in the previous section, could be due to a limitation of the backbone architecture. Predicting the next frame of a video with a contrastive loss requires learning invariances to every kind of transformation (or augmentation) that could happen from one frame to the next. For example, representing objects’ shapes requires invariance to objects’ motion, and representing objects’ motion requires invariance to objects’ shape. As suggested in [6], these two types of invariances often underlie the distinction between the dorsal and ventral representations in the brain. Therefore, one possibility is that ventral and dorsal-like representations compete for resources in the network, and ultimately, the ventral-like representations win out, possibly because they provide a greater overall reduction in loss. If this hypothesis is true, then a network with two separate pathways may be able to reduce the competition by assigning one pathway to be more ventral-like, and one more dorsal-like.

To test this hypothesis, we use the simplest extension of ResNet-1p: ResNet-2p, a ResNet architecture that is composed of two identical, parallel ResNets which split after the first layer and merge after their final layers (Fig. 3a; see also section 3.3). Each pathway of ResNet-2p has half the number of channels of ResNet-1p, keeping the total number of channels per layer equal between the two architectures. We choose this ResNet-2p architecture as it shares all of the features of ResNet-1p, but with two separate pathways that could potentially be assigned specialized functions. We then check whether ResNet-2p can learn both ventral- and dorsal-like representations. We compare the representations of all the layers of the two pathways in ResNet-2p (blue and red in the schematic in Fig. 3a) with all the visual areas. The results show that the representations along one of the pathways (the blue pathway in Fig. 3a-b, top) are more ventral-like, and the representations along the other pathway (the red pathway in Fig. 3a-b, bottom) are more dorsal-like. Therefore, the two pathways together can model both ventral and dorsal areas of mouse visual cortex. Compared to an untrained ANN, both ResNet-1p and ResNet-2p achieve high RSM similarity values for ventral areas (Figure 3c), with ResNet-2p showing a slight decrease in VISlm and VISpm compared to ResNet-1p. However, unlike ResNet-1p, which fails to model the dorsal areas, ResNet-2p outperforms ResNet-1p and the untrained ANN for area VISam by a wide margin (Fig. 3c). For VISal, maximum RSM similarity values for ResNet-1p and ResNet-2p are around the same level (Fig. 3c), though the similarity values of the red pathway representations do not drop as much throughout the network as the blue pathway representations do, indicating the general similarity of the red pathway to the dorsal areas. Examination of the RSM similarity between the two pathways shows that the representational geometries are quite different (Fig. S2). Moreover, when we compare the RSMs of the two pathways with those from ResNet-1p, we can see that ResNet-1p has representations that are a better match to the ventral-like pathway from ResNet-2p (Fig. S3). This supports the idea that, in the single pathway model, there is a competition between the two forms of representation that favours ventral-like functions, and which leads to specialized functions in the two-pathway architecture. Using architectures with more than two parallel pathways also does not improve representational similarities (Fig. S5).

The hierarchy index values for the ventral and dorsal areas are shown in Figure 3d. The hierarchy index for every area is calculated based on the model pathway that aligns best with that area in terms of representation similarity (blue pathway for VISlm, VISp, VISpm, and red pathway for VISam, VISal). The hierarchy indices are shown separately for the ventral and dorsal pathways. Similar to the results with ResNet-1p for the ventral pathway, VISp has a lower hierarchy index (0.25 ± 0) than VISlm (0.31 ± 0.05) and VISpm (0.29 ± 0.07). For the dorsal pathway, VISam has a larger hierarchy index than VISal (VISam: 0.82 ± 0.16 vs. VISal: 0.19 ± 0.15) which is also consistent with the anatomical/functional hierarchy index (Fig. 1c).

In terms of the predictive loss function, the performances of ResNet-1p and ResNet-2p are not significantly different (top-3 accuracy for ResNet-1p: 94.64 (0.68) and ResNet-2p: 93.472 (0.83)). However, it is worth noting that ResNet-2p has, in total, fewer parameters than ResNet-1p (ResNet-1p: 435k vs ResNet-2p: 285k). Therefore, considering the lower capacity of ResNet-2p and its similar predictive performance to ResNet-1p, we can conclude that the inductive bias of parallel architecture could help in predictive processing.

### 4.4 Supervised learning of action recognition with parallel-pathways is not sufficient for dorsal match

The inductive bias of an architecture with parallel pathways combined with the spatiotemporal dynamics of the video data could be enough to trigger the emergence of both ventral and dorsal-like representations, as defined in the previous section, regardless of the loss function used. To understand the role of the loss function, we trained the same ResNet-1p and ResNet-2p backbones with a supervised action classification loss on the UCF101 dataset (Fig. 4a). Based on the representation similarity plots in Figure 4b, both blue and red pathways seem to learn similar representations that are more ventral-like (also see Fig. S2). The maximum RSM similarity values in Figure 4c show that both ResNet-1p and ResNet-2p reach similarity values higher than the untrained ANN for ventral-like areas, but for the dorsal areas, and specifically VISam, neither architecture achieves good representational similarity. The low performance of the action classification models here could not be due to the low spatial resolution of the training videos as our comparisons with ANNs pretrianed with higher spatial resolution videos also show similar results (see supplementary section F).

**Figure 4:**
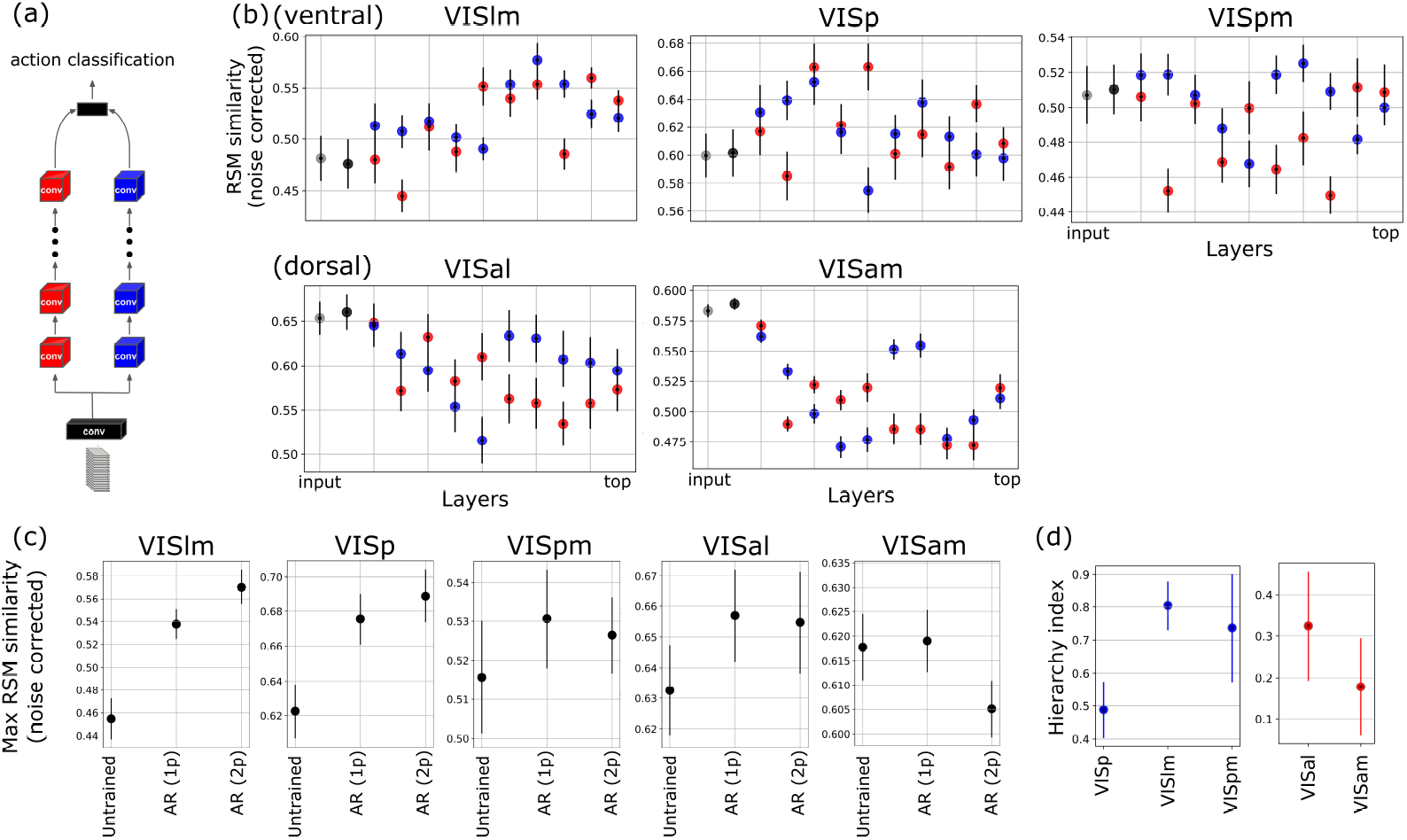
Representational Similarity Analysis between all the visual areas and the ANN trained with supervised action recognition loss function. (a) Representational similarity between all the layers of the ANN with two pathways (trained with action recognition objective) and the ventral-like (top: VISlm, VISp, VISpm) and the dorsal-like (bottom: VISal, VISam) areas. (b) The maximum representational similarity values between the ANN and the ventral and dorsal areas. AR (1p) and AR (2p) are ResNet-1p and ResNet-2p, respectively, both trained with action recognition loss function. (c) Hierarchy index of the ventral (left; in red) and dorsal (right; in blue) areas based on their fit to the ANN. Error bars represent bootstrapped standard deviation.

The hierarchy indices calculated using the ANN trained with an action classification objective (Fig. 4d), reproduce the CPC results (Fig. 3d and 2d) for ventral-like areas, and roughly match the anatomical/functional hierarchy index. However, the action classification model fails to predict the hierarchical organization of the dorsal areas, which is to be expected given the model’s poor alignment with these areas. Overall, these findings demonstrate the importance of the CPC loss function for learning both ventral and dorsal-like representations across the ResNet-2p architecture.

### 4.5 Functional specialization of the ventral and dorsal-like pathways in a CPC trained model

The ventral pathway is responsible for object and scene-based tasks [28], while the dorsal pathway is responsible for motion-based tasks [3, 47, 48, 54]. Based on our knowledge of the functional specialization of the two pathways in the real brain, we run a linear evaluation on the two pathways of ResNet-2p trained with CPC on two downstream tasks: (1) object categorization (CIFAR10 dataset) and (2) motion discrimination with random dot kinematograms (RDKs) (see Fig. 5a). In vision neuroscience, RDKs have been commonly used to characterise motion representation in the dorsal pathway [4]. We use this stimulus to evaluate the CPC-trained ResNet-2p (the red pathway in Fig. 3) on motion direction discrimination (four directions: up, down, left, right). Figure 5b shows the results for the two pathways, as well as for an untrained ResNet. As expected based on the comparisons with mouse visual areas, the ventral-like pathway is better in object categorization, while the dorsal-like pathway outperforms the ventral-like pathway in motion discrimination. Therefore, in addition to their better fits to neural data, the two pathways are indeed more ventral-like and dorsal-like according to their ability to support downstream tasks.

**Figure 5:**
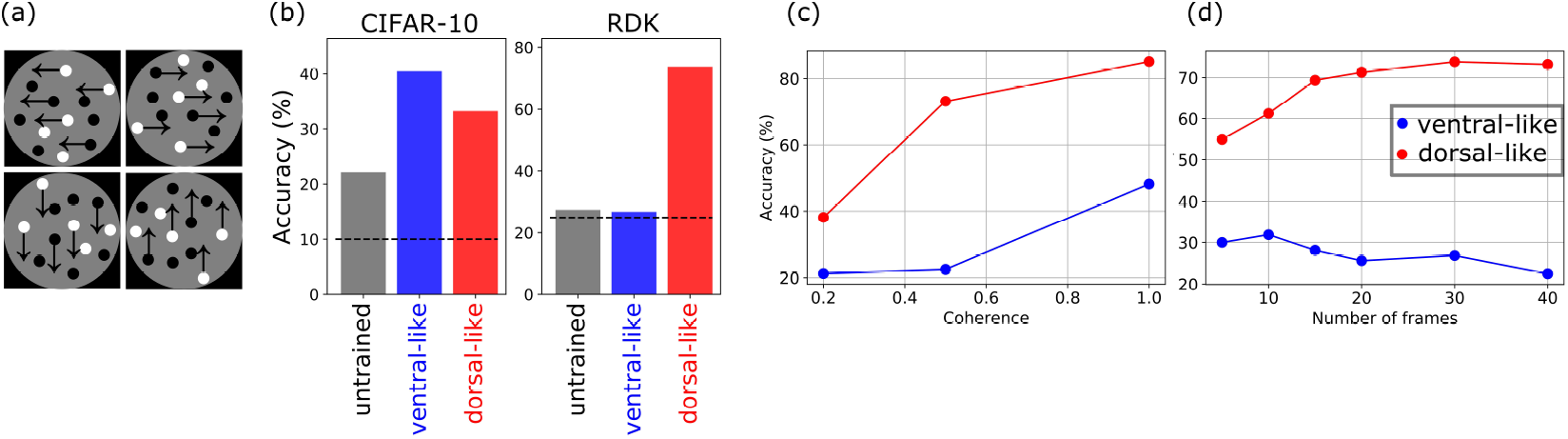
Random dots motion discrimination task. (a) Schematics of the RDK task. (b) The performance of the two pathways and an untrained ResNet on the object categorization (CIFAR-10) and motion discrimination (RDK) tasks. The black dashed lines show the chance levels for the two tasks. (c) The accuracy of the dorsal-like and the ventral-like paths for different levels of the random dots coherence. (d) The accuracy of the two pathways for different number of frames of the random dots stimulus.

As previously stated, decreasing the dots coherence (increasing spatial noise) makes the random-dots task more difficult. In real brains, dorsal areas can average out noise and extract motion signals by integrating motion over both time and space [46, 60]. It is thought that this is key to the importance of dorsal areas for RDK tasks. We measured ResNet-2p ventral-like and dorsal-like pathway performances for different coherence levels of random-dots (Fig. 5c). Ventral-like performance was dramatically reduced when the coherence was lowered from 100% to 50%. In contrast, the decrease in dorsal-like pathway performance was much smaller, demonstrating the dorsal-like pathway’s ability to spatially integrate motion, which is consistent with our expectations of dorsal areas. Indeed, the dorsal-like pathway can still achieve just under 40% accuracy even with 20% coherence. We also measured ventral-like and dorsal-like pathway performances for different numbers of frames of the random-dots stimuli at 50% coherence (Fig. 5d). The dorsal-like pathway’s performance improved significantly when the number of frames was increased, illustrating the ability of the dorsal-like pathway to integrate motion information over time. On the other hand, increasing the number of frames did not increase ventral-like pathway accuracy, but rather reduced it to around chance level. This shows that unlike the dorsal-like pathway, the ventral-like pathway of ResNet-2p does not integrate motion information over time.

## 5 Discussion

In this paper, we showed that self-supervised learning with CPC can produce representations that are more analogous to mouse visual cortex than either simple models or ANNs trained in a supervised manner. Furthermore, we showed that CPC applied to an architecture that has two parallel pathways can model both the ventral and dorsal areas of mouse visual cortex. The downstream tasks of object recognition and motion discrimination also support the ventral-like and dorsal-like representations of the two pathways in the model. Our experiments with supervised training on action classification indicated that the two-pathway architecture and video dataset are necessary but not sufficient for learning both types of representations, showing an interaction between the self-supervised objective function and the architecture. This finding shows that self-supervised predictive learning is a required component of our model for obtaining both ventral- and dorsal-like representations. Our observation that supervised action classification cannot generate dorsal representations is in contradiction with a previous fMRI study [21]. The different results of the two studies boil down to the different data modalities used in the two studies: two-photon calcium imaging in our study and fMRI in [21]. Capturing the responses of the dorsal areas elicited by natural videos with high temporal dynamics requires neuronal recordings with high temporal sampling rate. Therefore, we hypothesize that some aspects of dorsal representations of movement would not be reflected in the fMRI data, which could bias the fMRI-based analysis toward more static and lower temporal frequency features of the stimulus and neuronal responses (e.g. very slow-varying features; [61]).

Even though our results demonstrated the importance of self-supervised predictive learning for generating ventral- and dorsal-like representations, it is important to note that our conclusions are limited to the supervised loss functions that we included in our comparisons (i.e. object categorization and action recognition). We acknowledge that a different combination of more ecologically relevant supervised loss functions might be sufficient for learning ventral and dorsal representations in ANNs.

Learning representations of input data (images, videos, etc.) that are invariant to certain data augmentations (e.g. rotation, cropping, etc.) is a common goal of modern self-supervised learning methods. In models such as SimCLR [7] and BYOL [20], the augmentations are engineered for learning the most appropriate representations for downstream tasks. In CPC, however, the augmentations are inherent to the data being used. As noted above, predicting the next frame in a movie requires two different invariances: (1) invariance to motion, but selective for shape, and (2) invariance to shape, but selective for motion. Our results suggest that these two types of invariances are mutually exclusive, which can explain both the need for two separate pathways to get good matches to both ventral and dorsal areas and the inverse relationship between ventral-likeness and dorsal-likeness of the learned representation (see section D and Fig. S4 in supplementary materials). Thus, our results suggest that the functional specialization observed in the mammalian brain may be a natural consequence of a predictive objective applied to an architecture with two distinct pathways.

## 6 Limitations

One limitation of this work is the lack of comparisons with ANNs that are trained with other predictive loss functions, such as PredNet [38]. Other self-supervised video-based learning models (for example, see [14]) that do not optimize a predictive loss function could also be compared with CPC in terms of matching the representations of mouse visual cortex. Another limitation concerns the backbone architectures that we used in this study. Different parameters of the architectures (such as the number of residual blocks, the number of layers before the split and after the merger of the two pathways in in ResNet-2p, etc.) could be searched more thoroughly to determine the optimal setting for modeling mouse visual cortex. Furthermore, training ResNet-2p with CPC on a synthetic video dataset in which the motion and shape contents of the videos can be controlled could demonstrate more directly that the two pathways of ResNet-2p learn motion-invariant shape selectivity and shape-invariant motion selectivity, respectively.

## Broader Impact

ANNs can serve as a framework for understanding brains [50], as demonstrated here. This understanding would be based on finding the loss functions, architecture, and learning rules that best capture brain representations. Technologies that directly or indirectly interface with the brain, such as brain machine interfaces, can benefit from an *in silico* model of the brain. ANNs, if being used as such models, can facilitate designing and optimizing these technologies. However, the downside is that the ANN models of the brain are prone to the same limitations encountered by other ANNs, such as adversarial attack or implicit bias. These limitations can potentially leak into the applications in which these models would be used with human subjects.

## Acknowledgments and Disclosure of Funding

We thank Iris Jianghong Shi, Michael Buice, Stefan Mihalas, Eric Shea-Brown, and Bryan Tripp for helpful discussions. We also thank Pouya Bashivan and Alex Hernandez-Garcia for their suggestions on the manuscript. This work was supported by a NSERC (Discovery Grant: RGPIN-2020-05105; Discovery Accelerator Supplement: RGPAS-2020-00031), Healthy Brains, Healthy Lives (New Investigator Award: 2b-NISU-8; Innovative Ideas Grant: 1c-II-15), and CIFAR (Canada AI Chair; Learning in Machine and Brains Fellowship). CCP was funded by a CIHR grant (MOP-115178). This work was also funded by the Canada First Research Excellence Fund (CFREF Competition 2, 2015-2016) awarded to the Healthy Brains, Healthy Lives initiative at McGill University, through the Helmholtz International BigBrain Analytics and Learning Laboratory (HIBALL).

## Checklist

The checklist follows the references. Please read the checklist guidelines carefully for information on how to answer these questions. For each question, change the default **[TODO]** to [Yes], [No], or [N/A]. You are strongly encouraged to include a **justification to your answer**, either by referencing the appropriate section of your paper or providing a brief inline description. For example:

- Did you include the license to the code and datasets? [Yes] See Section **??**.
- Did you include the license to the code and datasets? [No] The code and the data are proprietary.
- Did you include the license to the code and datasets? [N/A]

Please do not modify the questions and only use the provided macros for your answers. Note that the Checklist section does not count towards the page limit. In your paper, please delete this instructions block and only keep the Checklist section heading above along with the questions/answers below.

1. For all authors…
  a. Do the main claims made in the abstract and introduction accurately reflect the paper’s contributions and scope? [Yes]
  b. Did you describe the limitations of your work? [Yes] See section 6
  c. Did you discuss any potential negative societal impacts of your work? [Yes]
  d. Have you read the ethics review guidelines and ensured that your paper conforms to them? [Yes]
2. If you are including theoretical results…
  a. Did you state the full set of assumptions of all theoretical results? [N/A]
  b. Did you include complete proofs of all theoretical results? [N/A]
3. If you ran experiments…
  a. Did you include the code, data, and instructions needed to reproduce the main experimental results (either in the supplemental material or as a URL)? [Yes] See the supplementary materials
  b. Did you specify all the training details (e.g., data splits, hyperparameters, how they were chosen)? [Yes] See section 3.3 and section A of supplementary materials
  c. Did you report error bars (e.g., with respect to the random seed after running experiments multiple times)? [Yes] All the values reported in the figures have error bars. The error bars represent the standard deviations estimated with bootstrapping on the neurons.
  d. Did you include the total amount of compute and the type of resources used (e.g., type of GPUs, internal cluster, or cloud provider)? [Yes] See section 3.3
4. If you are using existing assets (e.g., code, data, models) or curating/releasing new assets…
  a. If your work uses existing assets, did you cite the creators? [Yes] See section 3.3
  b. Did you mention the license of the assets? [N/A]
  c. Did you include any new assets either in the supplemental material or as a URL? [Yes] We included the codes and the checkpoint files for the pretrained models.
  d. Did you discuss whether and how consent was obtained from people whose data you’re using/curating? N/A]
  e. Did you discuss whether the data you are using/curating contains personally identifiable information or offensive content? [N/A]
5. If you used crowdsourcing or conducted research with human subjects…
  a. Did you include the full text of instructions given to participants and screenshots, if applicable? N/A]
  b. Did you describe any potential participant risks, with links to Institutional Review Board (IRB) approvals, if applicable? [N/A]
  c. Did you include the estimated hourly wage paid to participants and the total amount spent on participant compensation? [N/A]

## Supplementary Material

**Figure S1:**
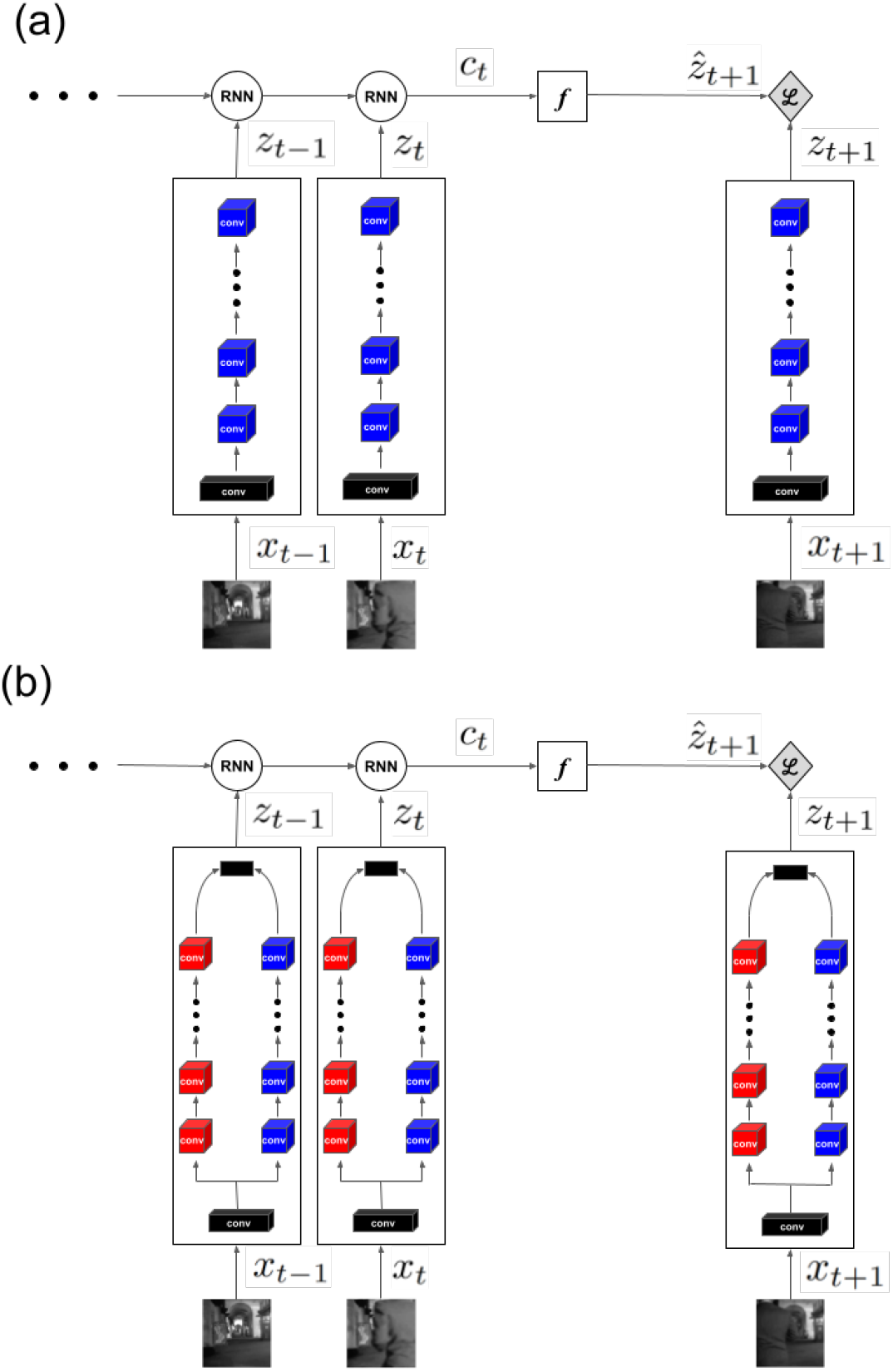
The schematic of Contrastive Predictive Coding model with (a) ResNet-1p, and (b) ResNet-2p backbone architectures. The present and past frames of the video (*x_t_, x*_*t*−1_,…) are given as input to the 3D ResNets. The output of the ResNets at each time point (*z_t_, z*_*t*−1_,…) are then passed to a recurrent neural network (RNN) which generates a context variable at time *t* (*c_t_*). The context variable, *c_t_*, is then used to predict the latent state of the next frame 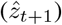 via a single layer MLP (*f*). The predicted next frame along with the latent representation of the correct next frame (positive sample, *z*_*t*+1_) and incorrect samples (negative examples, not shown here) are then fed to the loss function (see equation 1).

### A Supplementary methods

#### A.1 RSA noise ceiling

To estimate the noise ceiling of *RSM* similarity (the maximum *RSM* similarity we can reach for a given area), we randomly split the neuronal responses into two groups, and calculate two different *RSM*s based on each split. We then calculate the similarity between the *RSM* of the splits (as explained in section 3.2), and the similarity value gives us an estimate of the best possible match (i.e. the noise ceiling) for the recordings of that brain area. This process is repeated 100 times to also obtain a measure of variability for the estimated noise ceiling. The diagonal values of the matrix in Figure 1b shows the average noise ceiling values for every area.

#### A.2 Contrastive predictive coding

As noted in section 3.3, the CPC loss function relies on predicting the future latent representations of a video sequence, given its present and past representations (Fig. S1). Specifically, a block of *N* frames (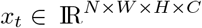; *W, H* are the spatial dimensions, *C* is the number of input channels) are passed through a backbone architecture, here a 3D convolutional neural network (CNN), and the CNN output (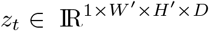; *W*′, *H*′ are the spatial dimensions, *D* is the number of output channels) is given to a recurrent neural network (RNN). The RNN aggregates the latent variables of S blocks of frames (*z*_*t*–*S*+1_,…, *z_t_*; here *S* = 5), and generates the context variable *c_t_* which is then used to predict the future *T* steps of the latent variables (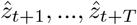; here *T* = 3). Importantly, the prediction is done in the latent space (*i.e*. the CNN output), and not in the pixel space. Specifically, CPC optimizes the following contrastive loss function:

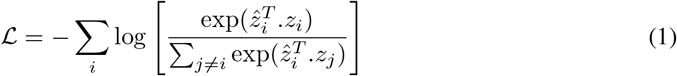

where, *i* and *j* denote the *i^th^* and *j^th^* time points. Minimizing the above contrastive loss function maximizes the similarity of the predicted latent variable 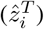 and the true future latent state (*z_i_*; AKA positive pairs), and minimizes its similarity with incorrect latent states (*z_j_* for *j* ≠ *i*; AKA negative pairs).

#### A.3 Other models

Two models are used as baselines: 3D Gabors, that were shown to be an acceptable model of the primary visual cortex in cats and primates [55, 29], and an untrained, randomly initialized 3D ResNet. The Gabor model comprised of 3D Gabor filters, spanning 8 motion directions (equally spaced between 0 and 315 degrees), 3 phases (−1, 0, 1), and 5 spatial scales. The randomly initialized model has its weights selected by matching the mean and standard deviation of synaptic weights in the trained models. These two baseline models are expected to be surpassed, in terms of alignment with brain representations, by any trained ANN that is using a relevant loss function for the visual system (meaning that the loss function captures something relevant about how either evolution or learning shape the brain). In addition to the baseline models, we also compare the CPC trained ANNs with ANNs that are trained with two other supervised loss functions: supervised object categorization (ImageNet dataset) and supervised action recognition (UCF101 dataset). We consider the ANNs trained with object categorization simply because they have been the standard in studies of the ventral pathway. We also use supervised action recognition (with UCF101 video dataset) in order to ensure that we have a fair comparison with CPC, which is being trained with dynamic videos. Importantly, both CPC and supervised action recognition use the UCF101 dataset, which makes the loss function the only difference between these two models. In order to show that the spatiotemporal dynamics of the video data is important, we also compared CPC with SimCLR, a self-supervised learning model trained on static images [7]. All the models are compared based on their representation alignment with different areas of mouse visual cortex (see section 3.2), and by using downstream tasks (see section 3.3).

#### A.4 Datasets

For training the deep ANNs, we use the UCF101 dataset. UCF101 is a dataset of 13320 short video segments from 101 action categories collected from YouTube [56]. We use this dataset for both self-supervised and supervised training of the ANNs. In the case of supervised training, the ANNs are trained to categorize actions in the videos. However, the UCF101 videos are not used for comparing representations between brain areas and the ANNs. For this, we use the same videos that the Allen Brain Observatory presented to the mice. Both datasets (UCF101 and Allen Brain Observatory videos) were normalized (with mean and standard deviation calculated across each dataset) and downsampled to 64 × 64 to account for the low spatial resolution of mouse retina.

#### A.5 Linear evaluation tasks

As noted in section 3.3, we examine the two pathways of our trained ResNet-2p on two downstream tasks: object categorization and motion discrimination, which are supported in the real brain by the ventral and dorsal pathways, respectively. For each downstream task, the weights of the trained pathways are frozen and a linear classifier is trained on the final convolutional layer of each pathway. We use the CIFAR-10 dataset for the object categorization task [36]. For the motion discrimination task, we use random-dot kinetograms (dot density = 2.5%, dot size = 2 *pixels*, speed = 5 *pixels/frame*). In every sample of the stimulus, a portion of dots move coherently in one of four directions (up, down, right, left), and the rest move completely randomly. The linear classifier is trained to detect the principal direction of motion. The models are evaluated on 3 motion coherence levels (20%, 50%, 100%) and 6 stimulus durations (5, 10, 15, 20, 30, 40 frames).

#### A.6 Code availability

The codes repository and the pretrained models are available at: https://ventral-dorsal-model.netlify.app/.

**Table S1:** An instantiation of the ResNet-1p and ResNet-2p architectures. The dimensions of kernels are shown by *T* × *S*^2^, *C* for temporal, spatial, and channel dimensions. The parameters of the two pathways of ResNet-2p are shown separately in red and blue colors. Output sizes are also denoted by *T* × *S*^2^ for temporal and spatial dimensions. The concatenate layer is only included in ResNet-2p where the two pathway outputs are concatenated along the channel dimensions. The sequence of res3 and res4 residual blocks repeats 4 times. In total, there are 10 residual blocks in both ResNet-1p and ResNet-2p.

| Stage | ResNet-1p | ResNet-2p | Output size |
| --- | --- | --- | --- |
| data layer | | | $5 \times 64^2$ |
| conv<sub>1</sub> | $5 \times 7^2, 256$<br>stride $1 \times 1^2$ | $5 \times 7^2, 256$<br>stride $1 \times 2^2$ | $5 \times 32^2$ |
| pool<sub>1</sub> | $1 \times 3^2$<br>stride $1 \times 2^2$ | $1 \times 3^2$<br>stride $1 \times 2^2$ | $5 \times 16^2$ |
| res<sub>1</sub> | $\begin{bmatrix} 1 \times 1^2, 256 \\ 1 \times 3^2, 32 \\ 1 \times 1^2, 128 \end{bmatrix}$ | $\begin{bmatrix} 1 \times 1^2, 128 \\ 1 \times 3^2, 16 \\ 1 \times 1^2, 64 \end{bmatrix}, \begin{bmatrix} 1 \times 1^2, 128 \\ 1 \times 3^2, 16 \\ 1 \times 1^2, 64 \end{bmatrix}$ | $5 \times 16^2$ |
| res<sub>2</sub> | $\begin{bmatrix} 3 \times 1^2, 128 \\ 1 \times 3^2, 32 \\ 1 \times 1^2, 128 \end{bmatrix}$ | $\begin{bmatrix} 3 \times 1^2, 64 \\ 1 \times 3^2, 16 \\ 1 \times 1^2, 64 \end{bmatrix}, \begin{bmatrix} 3 \times 1^2, 64 \\ 1 \times 3^2, 16 \\ 1 \times 1^2, 64 \end{bmatrix}$ | $5 \times 16^2$ |
| res<sub>3</sub> | $\begin{bmatrix} 1 \times 1^2, 128 \\ 1 \times 3^2, 32 \\ 1 \times 1^2, 128 \end{bmatrix}$ | $\begin{bmatrix} 1 \times 1^2, 64 \\ 1 \times 3^2, 16 \\ 1 \times 1^2, 64 \end{bmatrix}, \begin{bmatrix} 1 \times 1^2, 64 \\ 1 \times 3^2, 16 \\ 1 \times 1^2, 64 \end{bmatrix}$ | $5 \times 16^2$ |
| res<sub>4</sub> | $\begin{bmatrix} 3 \times 1^2, 128 \\ 1 \times 3^2, 32 \\ 1 \times 1^2, 128 \end{bmatrix}$ | $\begin{bmatrix} 3 \times 1^2, 64 \\ 1 \times 3^2, 16 \\ 1 \times 1^2, 64 \end{bmatrix}, \begin{bmatrix} 3 \times 1^2, 64 \\ 1 \times 3^2, 16 \\ 1 \times 1^2, 64 \end{bmatrix}$ | $5 \times 16^2$ |
| concatenate | — | | $5 \times 16^2$ |

### B Representational similarity between the two pathways of ResNet-2p

**Figure S2:**
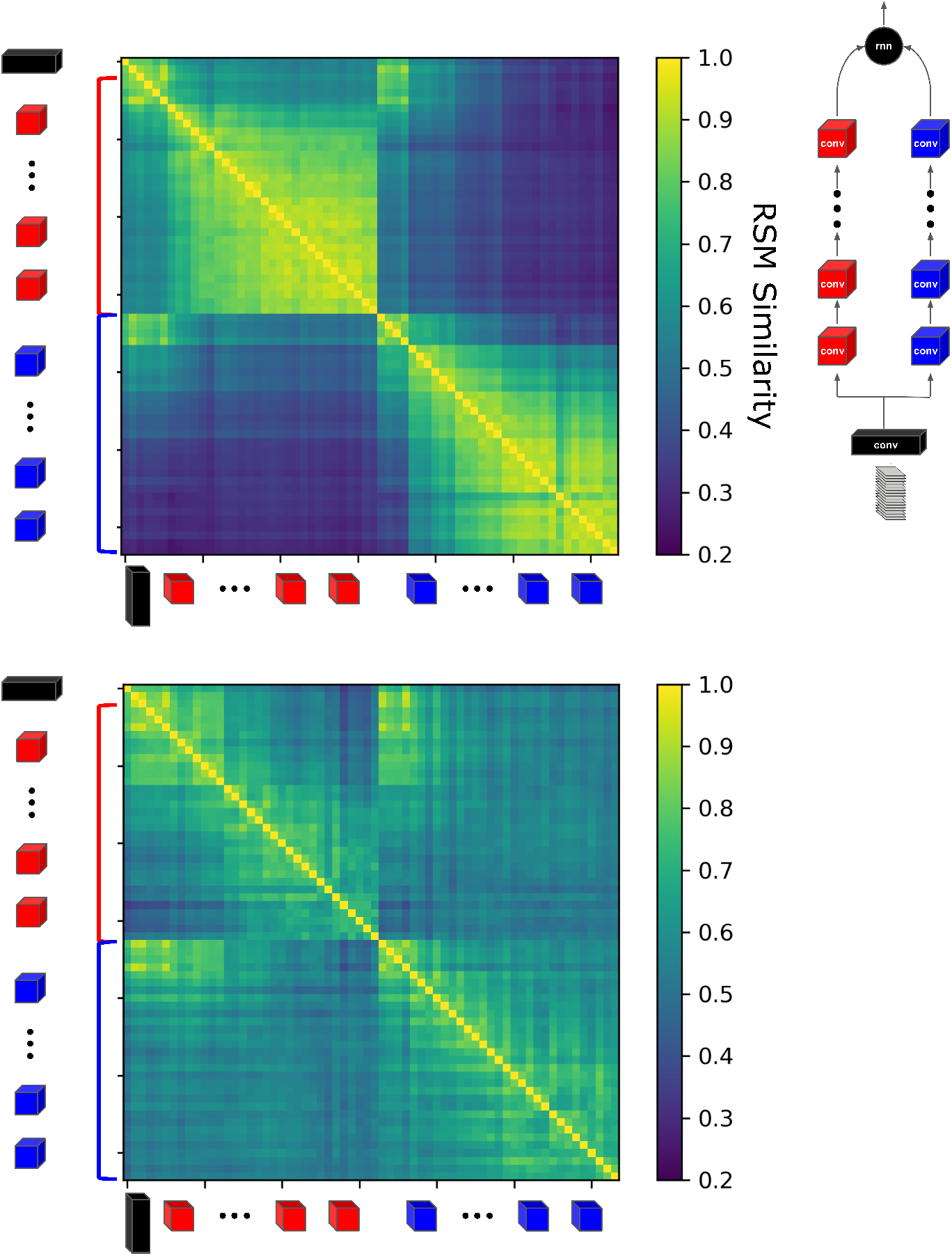
Representational similarity between and within the two pathways of ResNet-2p trained with CPC (top) and action recognition (bottom) loss functions.

### C Representational similarity between the two pathways of ResNet-2p and one pathway of ResNet-1p

**Figure S3:**
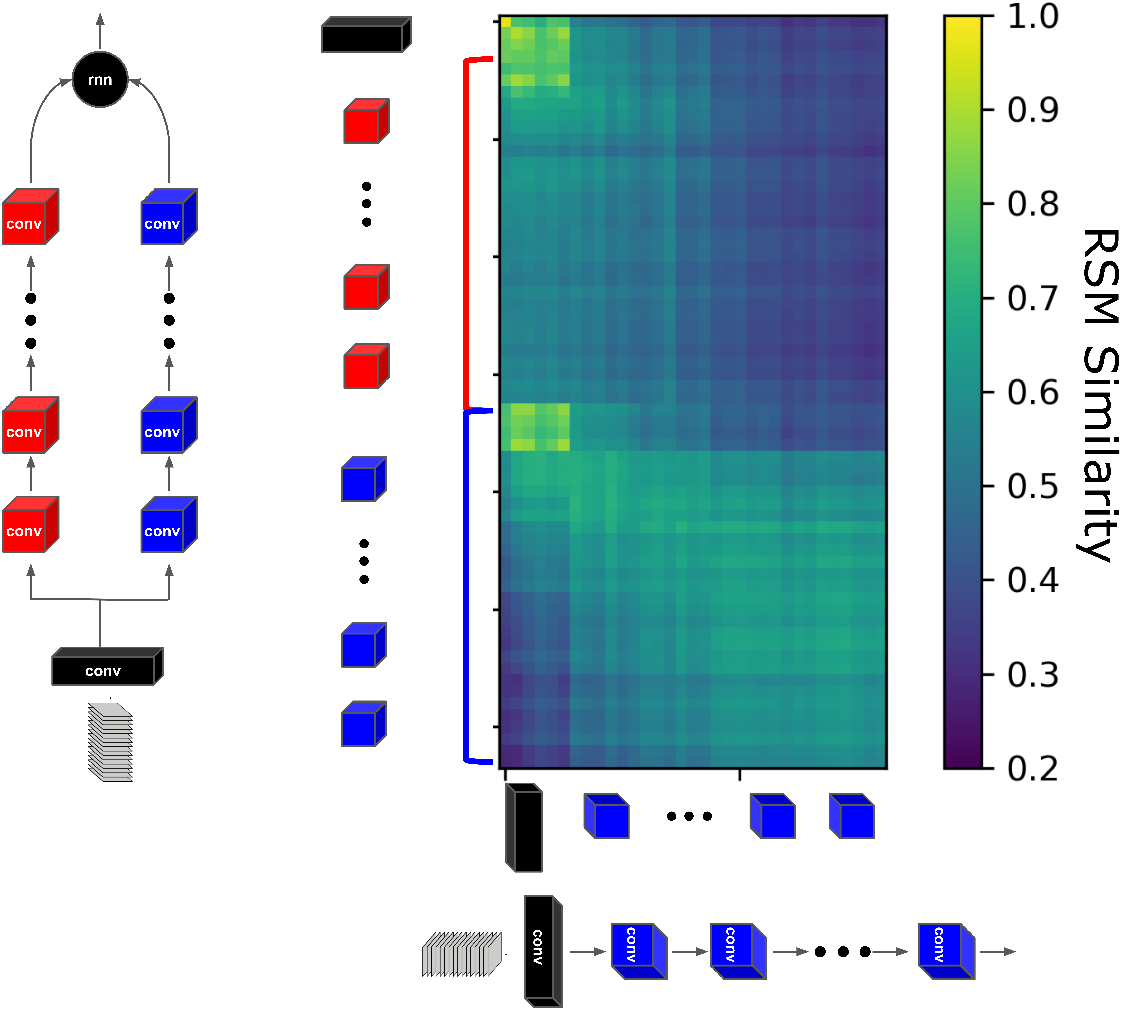
Representational similarity between ResNet-1p (columns in the figure) and ResNet-2p (rows in the figure) trained with CPC loss function. The blue and the red pathways in ResNet-2p schematic represent the ventral- and the dorsal-like pathways, respectively. As can be seen in the figure, the lower part of the rectangle has higher values than the upper part showing that ResNet-1p has more similar representations to the ventral-like pathway (blue in ResNet-2p schematic) than the dorsal-like pathway (red in ResNet-2p schematic).

### D Representational similarity during training

To gain an insight into the development of the dorsal- and ventral-like pathways, we quantify the similarity of each pathway in ResNet-1p and ResNet-2p to dorsal and ventral areas during training with CPC. For that purpose, we use the maximum representation similarity with VISam and VISlm (the most dorsal and ventral areas, respectively) as the dorsal-score and ventral-score, respectively. The scores are normalized between 0 and 1 and shown in Figure S2, for both ResNet-1p (left) and ResNet-2p (right). The results suggest that there is a competition between dorsal- and ventral-like representations in the one-pathway architecture. An increase in ventral score leads to a decrease in the dorsal score in ResNet-1p, which is reflected in the correlation value of *r* = −0:54 between the two scores. However, the second pathway in the ResNet-2p model decreases the anti-correlation between the two scores to *r* = −0:18. Indeed, by assigning the two representations to separate pathways in ResNet-2p, the two representations become partially independent. This result suggests that a two-pathway architecture can learn both ventral- and dorsal-like representations because it can partially decouple these two competing forms of representation.

For 10 random initialization seeds, we do get the dorsal/ventral split in 7 seeds (average D-score = 0.714, average V-score = 0.663). In 1 seed, both pathways are more dorsal-like (D-score = 0.688, V-score = 0.504) and in 2 seeds, more ventral-like (average D-score = 0.620, average V-score = 0.664). Also, the seeds with better dorsal/ventral splits across the two pathways reach higher average predictive performance at the end of training (top-3 accuracy with dorsal/ventral split: 93.88 (1.02), without dorsal/ventral: 91.80 (1.17)).

**Figure S4:**
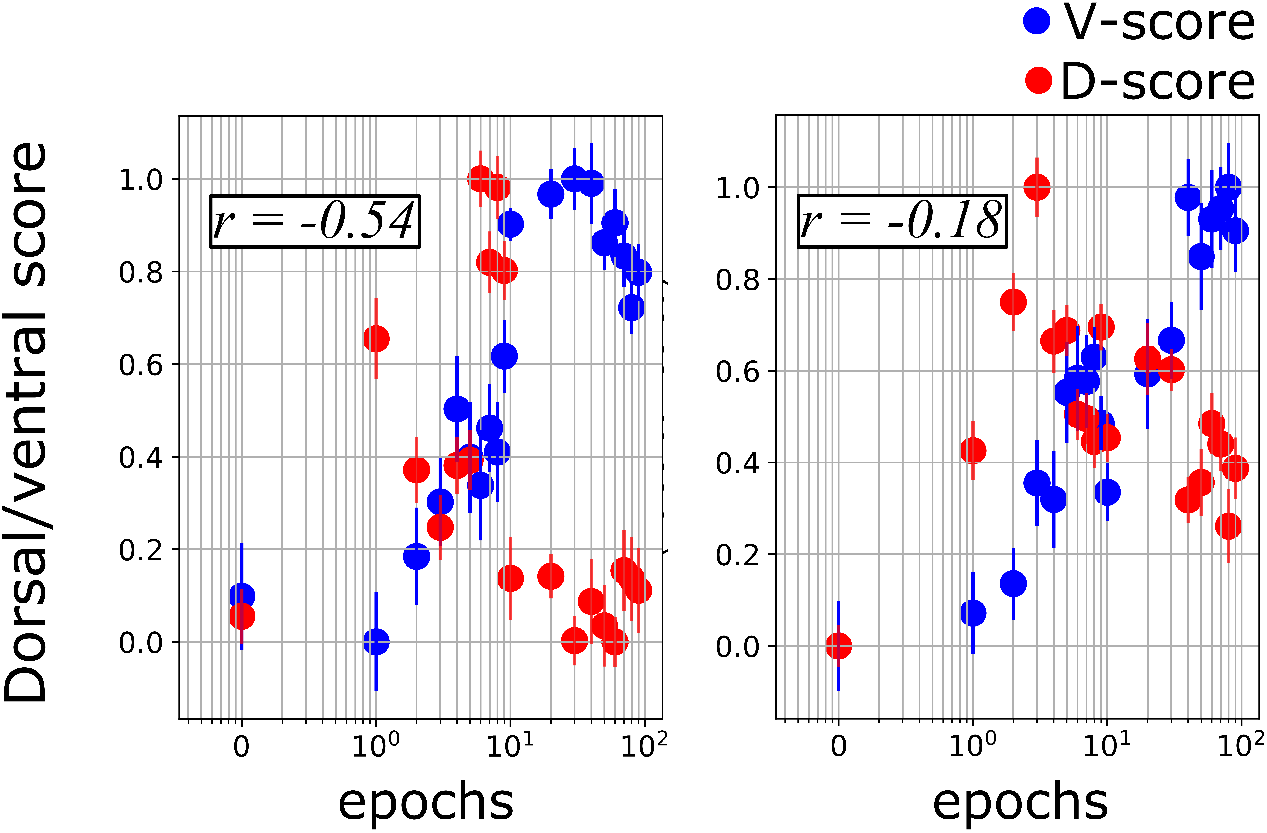
Ventral (red) and dorsal (blue) scores during training for the architectures with one (left) and two (right) pathways. *r* shows the correlation coefficient between the changes in dorsal and ventral scores during training.

**Figure S5:**
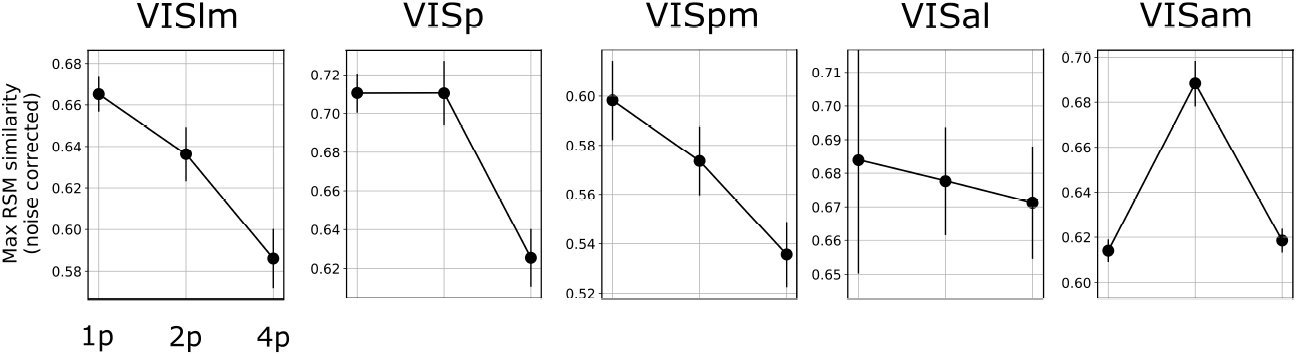
Maximum representational similarity for ResNet-1p, ResNet-2p, and ResNet-4p trained with CPC.For all the areas of mouse visual cortex, increasing the number of pathways from 2 to 4 decreased the representational similarity.

### E Dorsal-like specialization in the one-pathway model

Although the one-pathway architecture trained with CPC showed more similar representations with the ventral pathway in mouse visual cortex (Fig. 2), we can still ask explicitly if there is any learned dorsal-like specialization in ResNet-1p. To address this question, we take two approaches: (1) we measure the performance of ResNet-1p on the two downstream tasks, CIFAR-10 and RDK, as explained in section A.5, and (2) predicting the responses of the artificial neurons in ResNet-1p based on the responses of the neurons of the most ventral (VISlm) and the most dorsal (VISam) areas of mouse visual cortex. For (2), particularly, we take every artificial neuron in ResNet-1p, and fit a linear regression model to the neurons’ responses. We use 10-fold cross-validation to evaluate the linear fits. Neurons are labeled as dorsal-like if the dorsal neurons from VISam are better predictors of that artificial neuron’s responses than VISlm neurons.

The results of the two approaches are shown in Table S2 below. The linear evaluation results show that ResNet-1p is better than both ventral-like and dorsal-like pathways of ResNet-2p in object categorization, supporting the observation that ResNet-1p representations are more ventral-like. For the motion discrimination task, ResNet-1p is better than the ventral-like pathway of ResNet-2p, but worse than the dorsal-like pathway of ResNet-2p. This observation implies that ResNet-1p has some dorsal-like specialisation, but separating the two pathways in the architecture enhances dorsal-like specialisation in one of the pathways.

Consistent with the results of the downstream tasks, the linear regression approach also show that around 39% of ResNet-1p neurons are dorsal-like. When compared to the percentage of dorsal-like neurons in each pathway of ResNet-2p, we can see that using an architecture with parallel pathways enhances and isolates ventral and dorsal specializations in each pathway: the dorsal-like pathway has a significantly higher number of dorsal-like neurons compared to the ventral-like pathway and ResNet-1p.

**Table S2:**
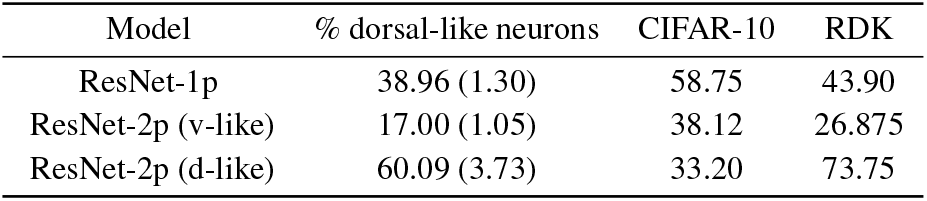
The percentage of dorsal-like neurons and the performance on downstream tasks (CIFAR-10 object categorization and RDK motion discrimination) for ResNet-1p and each pathway of ResNet-2p

### F Comparisons with pretrained high resolution action recognition models

The ANNs trained on supervised action classification, similar to the CPC-trained models, were trained with low spatial resolution videos (64 × 64; UCF101 dataset) to account for the low spatial resolution of mouse retina. The low resolution of the video data might affect the quality of the learned representations, especially in supervised learning. Therefore, in this section, we compare CPC with an ANN pretrained with higher resolution action classification datasets (112 × 112; Kinetics-400 dataset). We also compare the models with the SlowFast network: a two-stream architecture pretrained on action classification. However, unlike the two-pathway architecture used in our study, the two pathways of the SlowFast network are not identical. By choosing different stride sizes, kernel sizes, and input sampling rates, one pathway (the Slow pathway) was designed for learning representations of more static information, while the other pathway (the Fast pathway) was designed for capturing rapidly changing motion (see [13] for details).

The results of these comparisons are shown in Figure S6. First, we can see that the pretrained action classification model (AR (high res)) shows a lower representational similarity to all the brain areas compared to CPC, consistent with the results presented in Figure 4. Therefore, the lower performance of the supervised models, as reported in section 4.4, cannot be explained by the low resolution of the input videos that we used in the training.

Second, the SlowFast network performs at the same level as the action classification model for the two most ventral areas (VISlm and VISp) and the most dorsal area (VISam). Therefore, training an ANN with two parallel pathways on supervised action classification does not lead to learning ventral- and dorsal-like representations, consistent with the results in section 4.4 and Figure 4. However, the SlowFast network shows the highest representational similarity with two of the visual areas: VISpm (ventral) and VISal (dorsal). We can attribute the superior performance of the SlowFast network, in the case of these two areas, to the inductive biases used in this model. In particular, the temporal sampling rate of the input videos for training the Fast pathway of the SlowFast network was higher than the input sampling rate we used for training our models. While, in this study, we focused on the role of loss functions and number of pathways in the architecture, these results highlight the importance of other architectural hyperparameters in modeling different visual areas. Importantly, these results suggest that different visual areas may be best modeled with different architectural and input hyperparameters.

The pretrained action recognition model and the SlowFast network were both trained on Kinetics-400 which is a much larger dataset compared to UCF101 that we used in this study (650k samples in Kinetics-400 vs 13k samples in UCF101). Therefore, we should bear in mind that the training dataset is a confounding factor here when comparing the pretrainied models with CPC.

**Figure S6:**
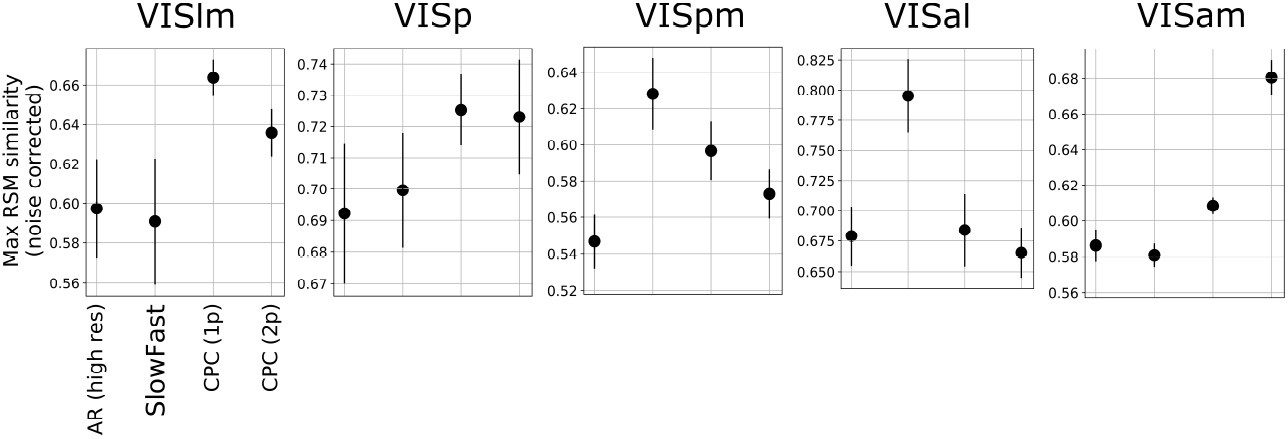
Maximum representational similarity for models trained on supervised action classification with high resolution videos (AR (high res)), the SlowFast network, and ResNet-1p and ResNet-2p trained with CPC.

